# Rapid decoding of neural information representation from ultra-fast functional magnetic resonance imaging signals

**DOI:** 10.1101/2025.07.21.665938

**Authors:** Yoichi Miyawaki, Kenshu Koiso, Daniel A Handwerker, Javier Gonzalez-Castillo, Laurentius Huber, Arman Khojandi, Yuhui Chai, Daniel Glen, Peter A Bandettini

## Abstract

High spatio-temporal resolution is crucial for neuroimaging techniques to improve our understanding of human brain function. While the fMRI signal is slow and shows a spread in latencies over space, the precision of hemodynamic response latency for each voxel is preserved and has been shown to be able to detect oscillatory hemodynamic changes approaching 1 Hz, suggesting its potential to reveal rapid neural dynamics. To examine how fast neural information can be derived from fMRI signals, we performed experiments that acquire high-field (7T) fMRI signals at an ultra-fast sampling rate (TR = 125 ms) from the visual cortex while participants observed naturalistic object stimuli. We applied multivariate pattern decoders to extract presented object-category information from the acquired signals at each sampling time after stimulus onset. Results showed that decoding accuracy rose above statistical significance less than 2 s after signal onset, faster than the peak latency of the hemodynamic response. The peak latency of the decoding accuracy was independent of variations in the hemodynamic latency of voxels used for decoding. The application of sparse decoders further revealed that rapid and accurate decoding was possible by pruning vein-rich voxels off from the multivariate voxel input to the decoders. These results suggest that a combination of ultra-fast sampling and multivariate decoding allows fast and temporally precise analysis of neural activity using fMRI signals.

**Significance statement:** Functional magnetic resonance imaging (fMRI) is the most successful method to evaluate human brain function at fine spatial scales but is thought to lack temporal resolution because of slow hemodynamics. We challenge this conventional notion by fast sampling of fMRI signals, combined with multivariate decoding to extract information content represented in the fMRI signals. Results showed that the information content can be extracted faster than the magnitude of hemodynamic response rises, independent of the large spatial variation of hemodynamic latencies across individual voxels, and even after removing venous voxels from decoders. Our method is thus effective at filtering out slow and variable hemodynamic components, leading to the extraction of rapid and temporally precise components reflecting neuronal activity in human fMRI signals.

## Introduction

The dynamic nature of the real world requires us to recognize our environment rapidly and accurately and take appropriate action. The mechanism for achieving such fast and precise information processing in the human brain has been an important question in neuroscience for many years. To answer this question, we need neuroimaging methods that have a high spatiotemporal resolution sufficient to analyze when and where the external information is represented in the brain. An example is the process of recognizing the various objects that make up our surrounding visual scene. The process is considered to happen sequentially and hierarchically in ventral high-level visual areas (1). Thus, we need a spatiotemporal resolution that can discern each subregion and the temporal difference in signal propagation between the subregions.

Functional magnetic resonance imaging (fMRI) has the high spatial resolution for such a requirement but is assumed not to have sufficient temporal resolution since it measures local blood flow changes. FMRI signals are typically modeled as the convolution of neural activity with the hemodynamic function, which has a large time constant and makes it difficult to retrieve dynamic information of the original underlying neural activity (2).

However, the above-mentioned shortcoming of fMRI is not necessarily correct when the characteristics of the fMRI signals are carefully examined. First, even if the time constant of the fMRI signal is large, the relative association between neural activity and blood flow signals is on the order of milliseconds. Menon et al. (1998) showed that the time difference between the temporally-modulated presentation of visual stimuli can be detected as the latency difference of fMRI signals with an accuracy of about 40 ms (3). More recently, Ekman et al. (2017) measured fMRI signals at intervals of 88 ms by limiting the number of slices to two, and succeeded in capturing fast transitions of brain activity corresponding to the perception of apparent motion (4). Lewis et al. (2019) also measured blood-oxygen-level dependent (BOLD) and cortical spinal fluid (CSF) signals at a sampling rate of 367-380 ms, and succeeded in analyzing the dynamics of brain activity during sleep (5). Second, the signal-to-noise ratio improves with static magnetic field strength, which enables fMRI signal measurement at even higher sampling rates. Lewis et al. (2016) has confirmed this prediction using a 7 T scanner, succeeding in the measurement of fMRI signals in response to visual stimuli changing at 0.75 Hz, which was not possible with a 3 T scanner (6). This evidence suggests the possibility that fast changes of neural activity might be captured by fMRI recordings obtained at a sufficiently high sampling rate with high SNR associated with high magnetic fields. Moreover, it also suggests the possibility of accessing fast dynamics of information representation in the human brain despite the slow and spatially variable hemodynamic-based fMRI signal.

From the viewpoint of capturing the dynamics of neural information processing from the fMRI signal, it is important to identify the timing when the information can be decoded. It is known that when decoding neural information represented in fMRI signals, voxels with a large amplitude are not always informative carriers (7, 8) since the large amplitude reflects signals from large veins, which pool blood from a wider area (9). Thus, to obtain fast dynamics of neural information processing, a multivoxel pattern decoding approach naturally selects for the most information-rich voxels that are likely smaller, more proximal venioles and capillaries.

To examine how fast neural information representation could be extracted from the fMRI signal, we implemented an object presentation paradigm using high-field (7 T), and a high sampling rate (TR = 125 ms). We retrieved information from the acquired signals on a volume-by-volume (i.e., time-resolved decoding) basis. In this study, we performed the following three analyses. First, we applied time-resolved decoding to examine how quickly visual stimulus information was able to be obtained from fast-sampled fMRI signals. Second, we evaluated the temporal relationship between decoding accuracy and hemodynamic response magnitude. Finally, we used sparse decoding (7) to identify locations in the brain from which the visual stimulus information could be predicted and subsequently to compare those to the spatial distribution of veins. These analyses demonstrate that neural information can be read out as early as the rising phase of the fMRI response, and that such rapid decoding arises from non-venous tissue, intrinsically filtering temporally smoothing large vessel effects by leveraging information content rather than signal magnitude.

## Results

### Fast fMRI signal acquisition at 8 Hz (TR = 125 ms) sampling

Each subject underwent two different types of scans. First, a single localizer scan (Figure 1A) that presented 20 s blocks of both scrambled and intact objects from eight different categories— namely “male”, “female”, “Siberian husky”, “toy poodle”, “airliner”, “fighter”, “jeep”, and “sports car”—was acquired to facilitate the subject-specific localization of higher-order visual cortex (HVC) involved in object processing. The scrambled and intact object blocks were presented alternatively in this order (each block was presented four times) followed by an extra scrambled object block at the end. Durations of the first and the last scrambled block were set as 10 s to equalize the total presentation duration for each condition. Thus, the total duration of the functional localizer session was 320 s. Second, an event-related scan (to which we refer as the decoding session throughout the manuscript; Figure 1.B) was obtained as we presented the same object category images (although not the same image exemplars) for 500 ms, followed by a rest period of randomly varied duration (uniformly random-sampled from a range of 8.5-13.5 s at a step of TR). Randomization of inter-stimuli interval was performed to avoid participants predicting the next stimulus presentation timing and to improve statistical independence of trial-based hemodynamic responses (Figure 1B). This combination of stimulus and rest constituted a single trial, and the time course of fMRI signals and decoding performance was analyzed from the stimulus onset to the end of each trial as typically done in an event-related paradigm. The duration of the rest period is shorter than the duration of the typical HRF, leading to the overlap with the next trial. However, such overlaps appear not to influence the accuracy of the decoding analysis (8, 10). Each run of the decoding session consisted of 32 trials preceded by a 12-s extra rest period at the beginning of the run. The total duration of each run was kept at 380 s by adjusting the durations of randomly chosen rest periods. Further details of the experimental design are described in Materials and Methods section.

**Figure 1:**
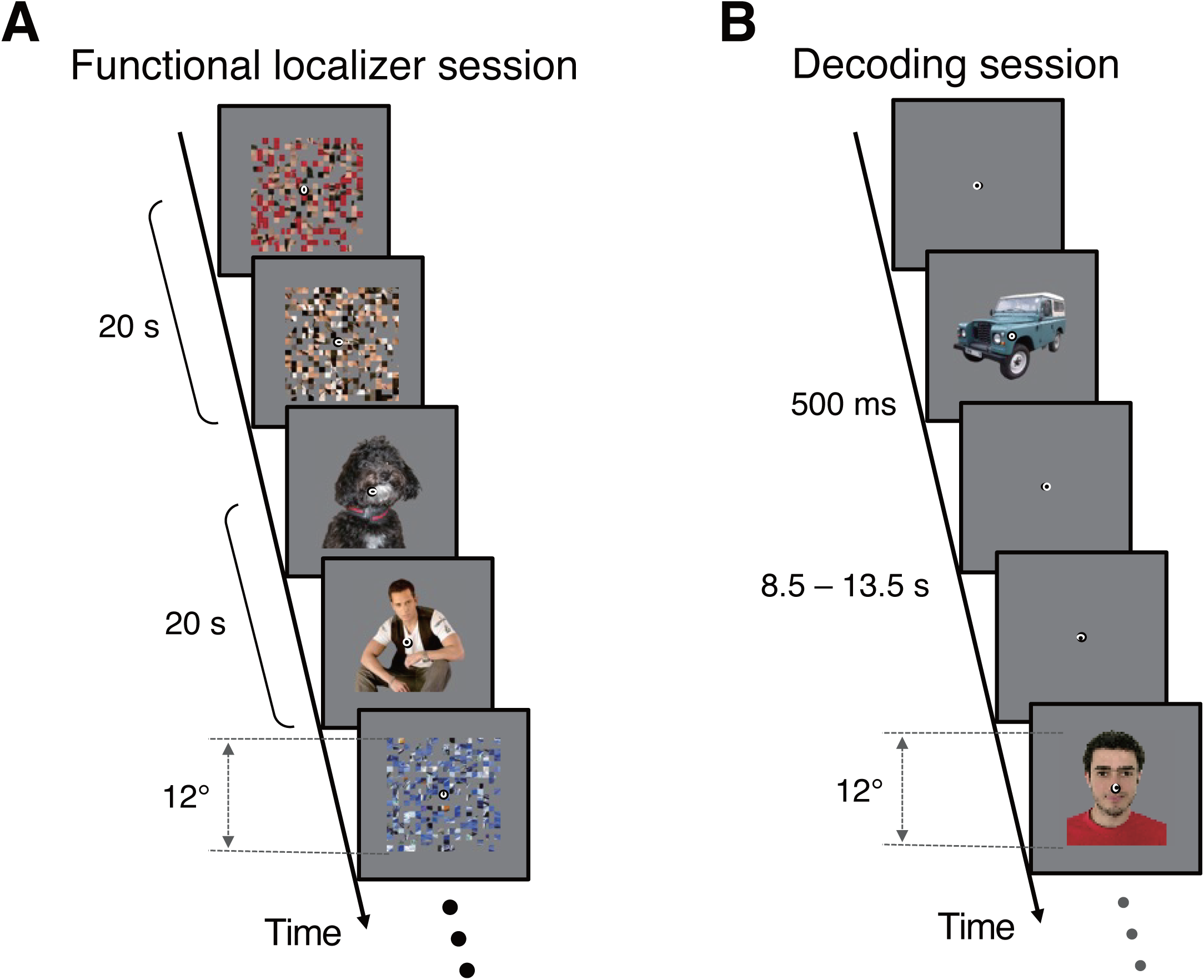
Experimental design. The experiment consisted of the functional localizer session and the decoding session. (A) Functional localizer session. Scrambled and intact object images (12 x 12 deg) were flashed in the 20-s stimulus block. The scrambled and intact blocks were presented alternatively to identify cortical areas that showed high contrast in fMRI signals between them. (B) Decoding session. Intact object images were presented for 500 ms followed by the rest period presenting a homogenous gray background image to examine the time course of hemodynamic response and decoding performance after the stimulus onset. The duration of the rest period was randomly chosen between 8.5 and 13.5 s. The same object categories but different exemplars were used across runs in the decoding session. Participants were instructed to press the button when the fixation color changed for 250 ms (the interval of the color changes was randomly sampled in a range of 1-9 s). This task was used in both the functional localizer and decoding sessions.

To test how fast stimulus information can be extracted from fMRI signals, we acquired fMRI data using a fast temporal sampling (TR = 125 ms) on a 7 T scanner. The spatial coverage was restricted to nine slices covering the ventral higher visual cortex (see Supplementary Figure 1 for the field of view for scanning) that responds to object categories and the primary visual cortex for comparisons.

Figure 2 shows a representative activation map from a single subject (subject S2) for the functional localizer session. This map suggests the presence of sufficient SNR on our data regardless of the short TR, given that data from only a single run was able to localize the HVC area, consistent with a previous study showing that short TR can provide gains in statistical power (11). Similar results were obtained for all other subjects (see Supplementary Figure 2A for all other subjects).

**Figure 2:**
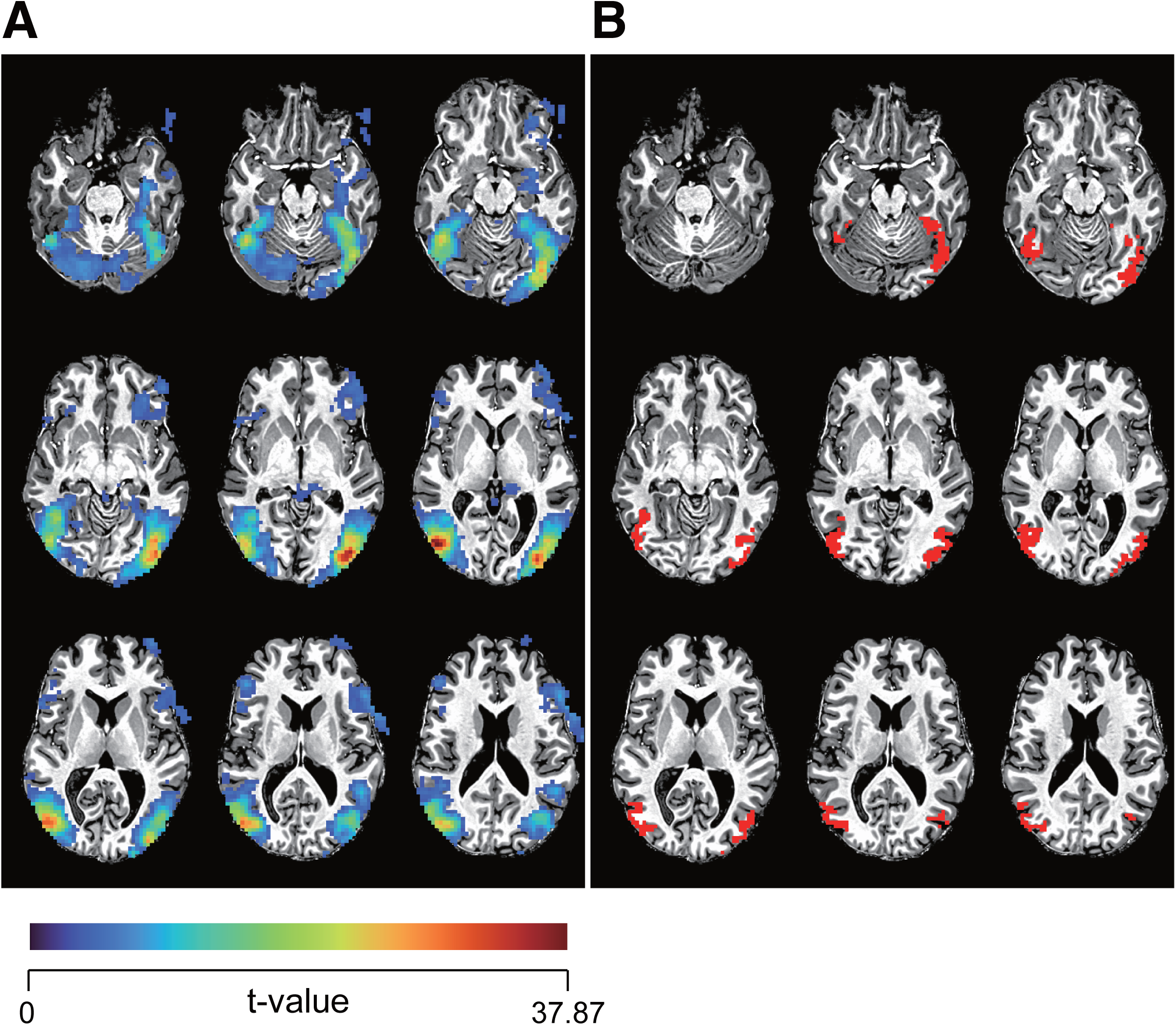
Functional localization with short TR signal acquisition. (A) Statistical t-value maps of fMRI signal contrast between intact–scramble object images. Results from a single participant (S2) are displayed as representative examples. Each functional slice is overlaid on the anatomical image. Voxels with one-sided p < 0.001 (uncorrected t-test) and clustering with more than 40 neighbors (voxel faces touch each other) are only shown with color. The colorbar is shown at the bottom and scaled between 0 and the maximum t-value (37.87 for this participant’s case). The in-plane voxel size is 3 x 3 mm and the slice thickness is 5 mm (see also Methods; each image is thus 5 mm apart in the inferior-superior direction). (B) The location of the top 400 t-value voxels from the individual gray matter (red colored) of the same participant (S2). These voxels were used for the decoding analysis. All other participants’ results are shown in Supplementary Figure 2 for panel (A) and Supplementary Figure 3 for panel (B), respectively.

### Rapid decoding from fast fMRI signals precedes fMRI signal evolution

We then identified the HVC area from the results of the functional localizer session as a region-of-interest (ROI) (see Materials and Methods for details to define the region of interests; voxels selected as input to the decoders are shown in Figure 2B and Supplementary Figure 2) and examined when it was possible to predict, or decode, presented object categories from fMRI signals in the HVC area. A decoder was trained and tested at each time point (i.e., TR or volume) sequentially, and the temporal change of the decoding performance was evaluated from the onset of the visual stimulus presentation to the end of each trial (“time-resolved decoding”). To remove the high-frequency variability in fMRI signals due to the short TR, we averaged the fMRI signals over three volumes before and after each time point (i.e., ±375 ms time window) and the temporally averaged fMRI signals were input to each decoder. Although eight object categories were used in the experiment (see Materials and Methods), here we present the decoding results for classifying two superordinate categories: “animal” and “vehicle”, each of which consisted of four subordinate categories (“male”, “female”, “Siberian husky”, and “toy poodle” for the “animal” category, and “airliner”, “fighter”, “jeep”, and “sports car” for the “vehicle” category). The results indicate that the presented object categories can be predicted approximately 2 s after the stimulus onset (red trace on Figure 3A for a single participant (S2) and Figure 3B for the mean across participants).

**Figure 3:**
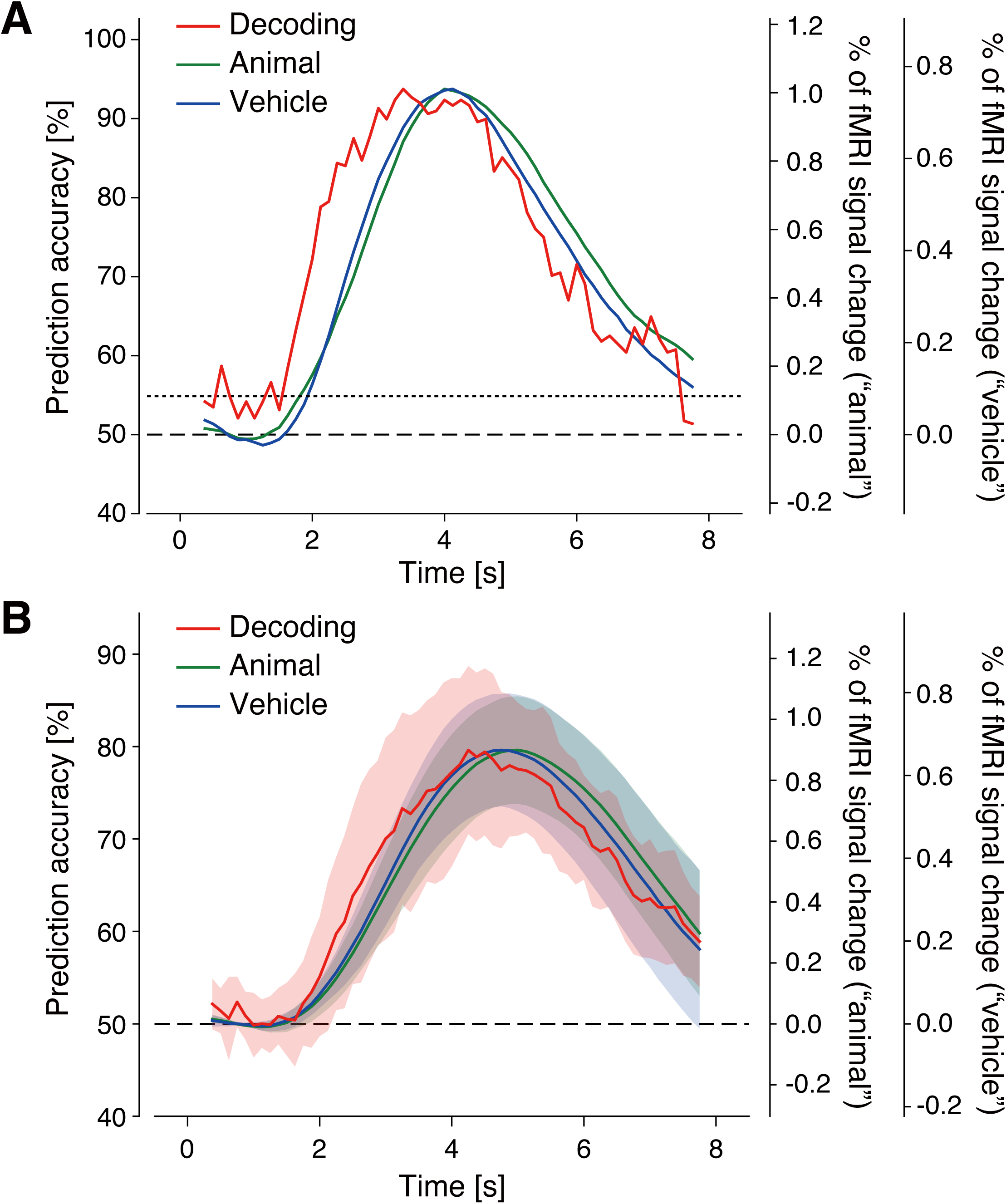
Rapid decoding of visual object categories. (A) The time courses of decoding performance and fMRI signals. The red line indicates the performance of the decoder in predicting presented object categories (“animal” or “vehicle”), which is plotted against the left axis. The green and blue lines are the percentages of fMRI signal changes (averaged over trials and voxels in the ROI shown in Figure 2B) from the baseline period (4 s) at the beginning of each run for each stimulus category, and are plotted against the right axes. Each vertical axis is rescaled so that (i) 50% prediction accuracy aligns with 0% fMRI signal change and (ii) the maximum prediction accuracy matches the maximum fMRI signal change, to visualize the difference in speed to reach their peaks. The dashed line represents the chance level for prediction accuracy (50%) and the baseline for fMRI signal change (0%). The dotted line represents the statistical significance level for the prediction accuracy (one-sided binomial test p < 0.05). Shown are examples from a single participant (S2) who showed the largest negative cross-correlation lag (decoding performance led ROI-averaged fMRI signals) among all participants. (B) The time courses of decoding performance and fMRI signals averaged across participants. Shaded areas indicate ±1 s.d. across participants. Each vertical axis is rescaled as in panel (A). The dashed line represents the chance level for prediction accuracy (50%) and the baseline for fMRI signal change (0%).

To examine the information source of the rapid decoding, we compared the time course of the decoding performance with the hemodynamic response function (HRF) in the ROI (HVC in this case) evoked by each of the object categories. The HRF was estimated by averaging the fMRI signals from all voxels used as input for the decoder (i.e., ROI-averaged HRF). As Figure 3 shows, the decoding performance rose faster than the HRFs of both the “animal” and “vehicle” categories. Since amplitude differences in the time course of the HRFs for each category might happen in their early phase, it is possible that such differences contributed to the success of decoding in early time points. To rule out this possibility, we used the averaged fMRI signal over voxels in the ROI while ignoring multivoxel pattern information in the ROI and examined how fast and accurately object category information could be decoded. The results showed that the ROI-averaged HRF was useful for object category decoding, but its performance was just above the chance level and significantly lower than the original performance using multivoxel pattern information (Figure 4). These results suggest that the rapid and accurate decoding observed in Figure 3 is based on the higher spatial resolution information used in multivariate patterns of fMRI signals within the HVC, not based on the difference in the mean magnitude of fMRI signals between the object categories.

**Figure 4:**
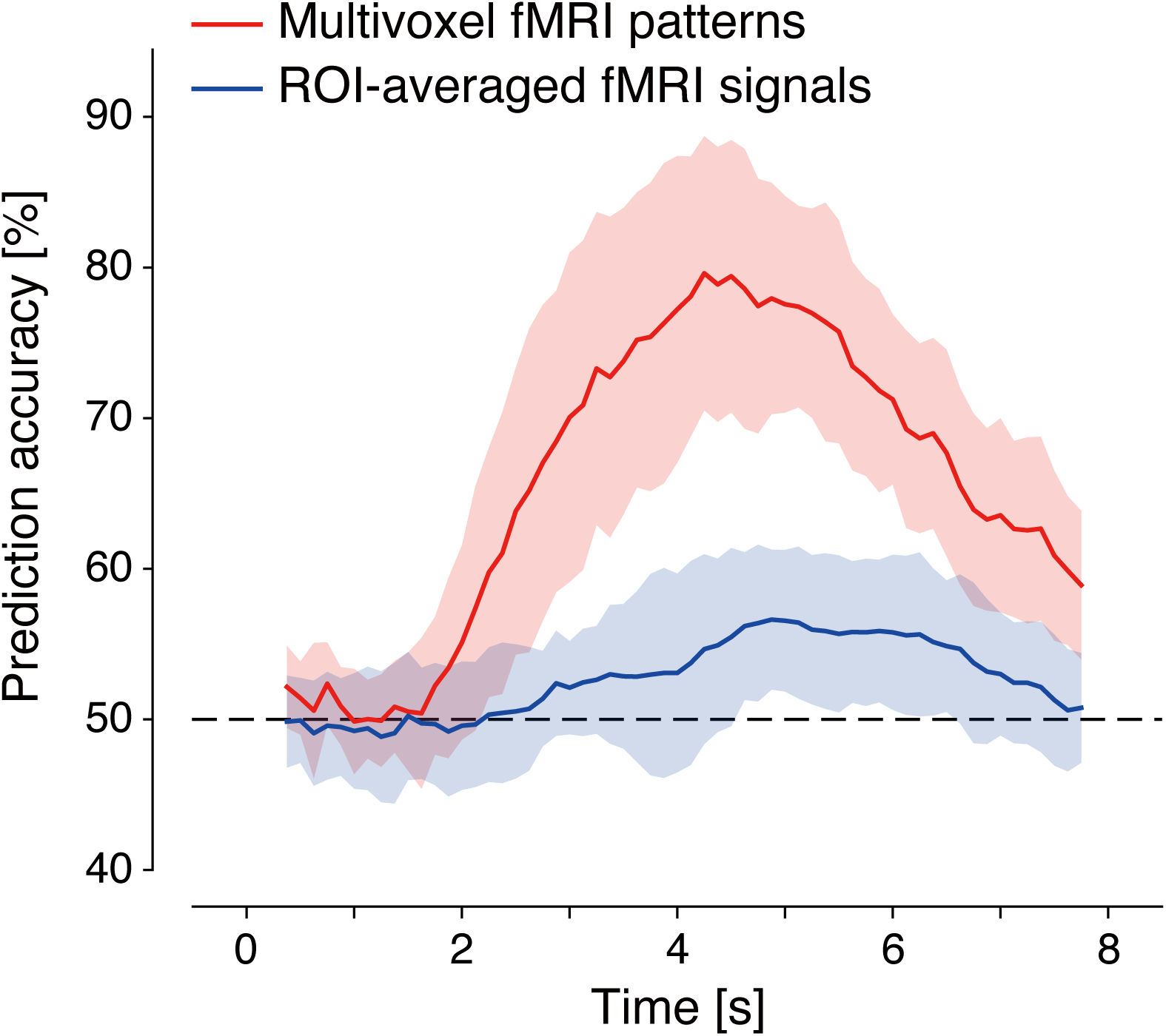
Comparison of decoding performance between multivoxel fMRI patterns and ROI-averaged fMRI signals. The red line indicates the performance of the decoder using multivoxel patterns of fMRI signals as input to the decoders (identical to the red line in Figure 3B). The blue line indicates the performance of the decoder using ROI-averaged fMRI signals as input to the decoders (mean over participants; shade, s.d.). The dashed line is the chance level.

### Temporal dissociation of decoding performance and hemodynamic response

To further investigate the information source of the rapid decoding, we next examined the latency difference in the multiple voxels in the ROI. Each individual voxel signal would have a different temporal evolution, revealing different temporal characteristics from the ROI-averaged HRF shown in Figure 3. Previous studies demonstrated that the latency of fMRI signals at each individual voxel varies widely, up to 4-5 s (2). We also confirmed that the peak latency of fMRI signals for each individual voxel varies approximately from 2-6 s after the stimulus onset (Figure 5A (“animal” category) and 5B (“vehicle” category) for a representative participant (S2), the voxels were sorted according to their peak latency), revealing the broad temporal distribution within the ROI. The lag of ROI-averaged HRF and the corresponding decoding performance are shown in Figure 5C (“animal” category) and 5D (“vehicle” category). The cross-correlation between decoding performance and individual-voxel HRFs shows that the distribution of the lag significantly shifts to the negative direction, indicating that the temporal evolution of decoding performance was faster than the latency of the fMRI signal for a large number of voxels, though there were individual differences in how much decoding performance precedes HRF latency (Supplementary Figure 4 showing that 2 out of 13 showed decoding was slower on average). On the other hand, the cross-correlation lag distribution also shows that there are voxels with positive or zero values, indicating that the temporal evolution of fMRI signal magnitude preceded/coincided with that of decoding performance. Based on these results, we hypothesize that the voxels with the short HRF latencies contribute most heavily as the information source for rapid decoding while those with the long latencies do not contain information necessary for the decoding.

**Figure 5:**
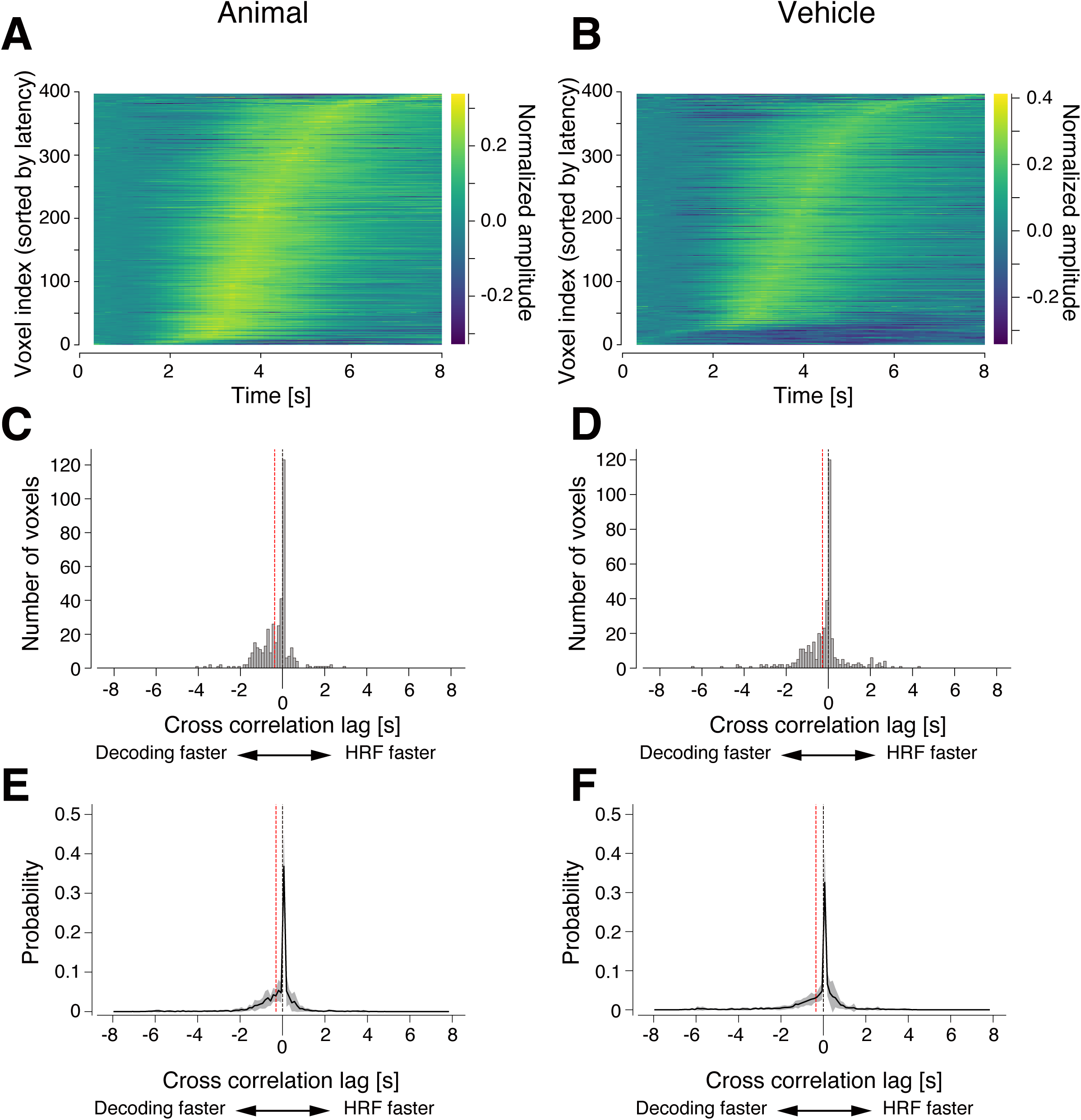
Temporal precedence of decoding performance over single voxel HRFs. (A, B) Latency distribution of single voxel HRFs for “animal” (A) and “vehicle” (B) category stimulus presentation. Voxels used for the decoding analysis are sorted by positive peak latency of trial-averaged fMRI signals in ascending order from bottom (shortest latency) to top (longest latency). The amplitude is normalized by the L2-norm of the trial-averaged fMRI signals over time and displayed as a pseudocolor (the corresponding color bar is on the right side of the plot). (C, D) Histograms of cross-correlation lags between the time course of decoding performance and the HRF of individual voxels. The black and red dashed lines indicate the zero lag and the average lags across voxels, respectively. Panels A to D are from a single participant (S2). The mean lags are significantly less than 0 for both categories (t-test, p < 0.05). (E, F) The probability distribution of the cross-correlation lag averaged over participants (shade, s.d.) for each category. Because the number of voxels actually used for decoding differed slightly between participants due to the outlier voxel rejection process, the number of voxels in the cross-correlation histogram was normalized by the total number of voxels and converted to the probability distribution before calculating the mean across participants. Black and red dashed lines indicate the zero lag and the mean lags across participants, respectively. The mean lags are significantly less than 0 (t-test, p < 0.05) for both categories.

To test this hypothesis, we divided voxels within the ROI into four groups based on their peak latency of voxel-wise HRFs and performed the same decoding analysis using only the voxels contained within each group (Figure 6A illustrating the procedure). If the hypothesis is correct, the group with shorter HRF latencies would result in faster decoding. However, the results indicate that decoding was possible from all latency groups, with little difference in the temporal evolution of prediction accuracy as well as the peak timing, though the maximum decoding accuracy tended to be slightly higher in the group with shorter latencies (Figure 6B). The peak latency of decoding in the short-latency group was almost identical to the HRF latency, but the divergence between decoding peak latency and HRF peak latency became more pronounced for the long-latency groups. This result indicates that information about the visual stimuli can be extracted from the multivoxel fMRI signal patterns at a similar time, weakly dependent on the temporal profile of contributing fMRI signal. It is known that within any voxel, the BOLD fMRI signal contains components related to small and large vessels. So it seems that the latency of the BOLD signal is dominated by large vessel effects whereas the small vessel component contributes to the early phase of the signals that provide sufficient decoding information. In summary, the information source of rapid decoding is the early fMRI signal component closely related to fast capillary oxygenation changes with neural activity, whereas the late large-vessel components contribute little to the prediction.

**Figure 6:**
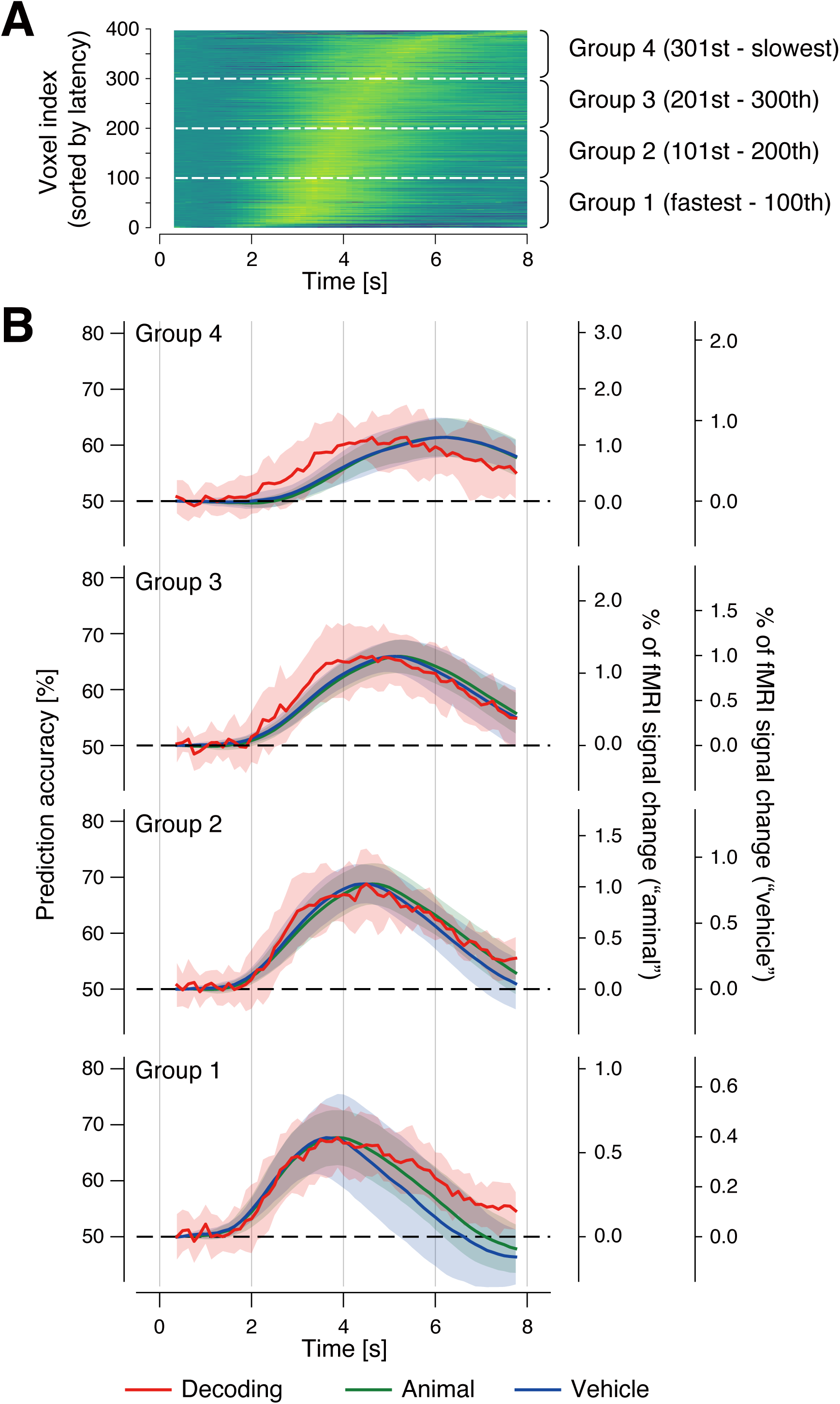
Temporal dissociation of decoding performance and hemodynamic response. (A) Schematic of voxel selection based on the latency of the fMRI signal of each voxel. Voxels input to the decoders are divided into four groups of approximately 100 voxels each according to the latency of the fMRI signal. (B) Comparisons between the time course of decoding performance and HRFs for each latency group. Results are presented using the same conventions as in Figure 3B for each latency group.

### Non-venous voxel selection by sparse decoders for rapid decoding

Voxels overlapping with venous regions are more likely to yield BOLD signals that strongly reflect large vessel characteristics, whereas those on the cortical tissue might reflect neural activity more strongly (2). If the results presented in Figure 6 show that the decoder automatically weighs the fMRI signals related to neural activity more tightly, then the voxels from which the decoder extracts information are expected to be more prevalent in non-venous regions than venous regions. To validate this hypothesis in a data-driven approach, we further acquired a susceptibility-weighted image (SWI) that excels at depicting venous regions as differences in the luminance of images (veins are illustrated as dark regions) and examined whether the voxels used by decoders overlap with the venous regions. A representative example of SWI from a single participant is shown in Figure 7A, which clearly illustrates veins as darker regions. We then computationally identified venous (darker) and non-venous (brighter) voxels by clustering based on the distribution of corresponding luminance values (for details, see Materials and Methods).

**Figure 7:**
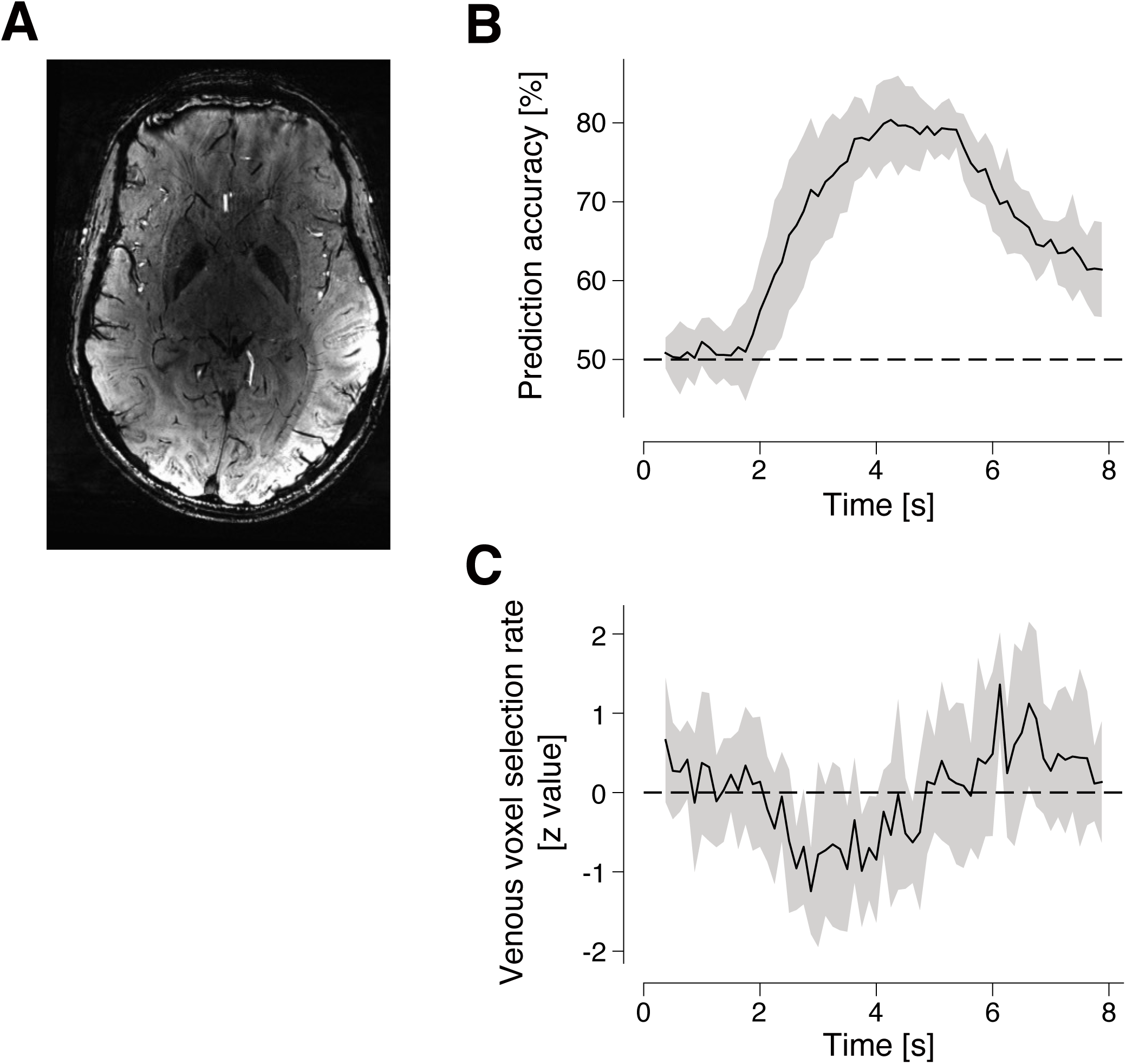
Non-venous voxel selection by sparse decoders for rapid decoding. (A) A susceptibility-weighted image for a representative participant (S2). Darker thin structures correspond to areas containing large veins. (B) The time course of prediction accuracy of time-resolved decoding using sparse decoders averaged across participants (shade, s.d.). The dashed line indicates the chance level. (C) The time course of the venous-voxel selection rate by sparse decoders averaged across participants (shade, s.d.). The venous-voxel selection rate is converted to a z-value based on the mean and the standard deviation calculated over the duration of the trial. The dashed line indicates the mean value. Thus the negative values indicate that the venous voxels are less selected.

To explicitly identify which voxels were used by the decoder to extract visual stimulus information, we employed a sparse logistic regression (SLR) model that sparsely selects informative voxels through the learning of the decoder and prunes off the uninformative voxels (7). More specifically, SLR promotes the linear weights of the unnecessary voxels going to zero, and only a small number of voxels remain with non-zero weights after learning (for details, see Materials and Methods). The decoding performance using SLR is shown in Figure 7B. The time course of prediction accuracy was generally the same as that obtained by the SVM model. It rose beyond the chance level around 2 s and reached a peak around 4 to 5 s after the stimulus onset. The peak decoding performance exceeded 80%, indicating that the decoders were successfully trained to achieve rapid decoding with accurate prediction using the SLR model while selecting informative voxels sparsely.

Then, we identified the locations of the selected voxels and quantified how much the selected voxels overlapped with venous locations at each time point after the stimulus onset. Figure 7C shows the time course of the venous voxel selection rate (the number of venous voxels among the selected voxels divided by the number of the selected voxels). Because the absolute values of the venous voxel selection rate varied largely across participants, we converted each participant’s values into z-scores and then averaged the z-scores across participants (see Materials and Methods section for details). The results reveal that the venous voxel selection rate significantly decreased at times when decoding performance rose before it reached its peak.

These results suggest that rapid decoding is likely achieved by extracting visual stimulus information selectively from non-venous voxels that reflect neural activity more strongly at short latency. In other words, the decoder naturally filtered out the non-neuronal, vascular-related characteristics from the fMRI signals, successfully extracting and analyzing signal components related to neural information processing.

## Discussion

We have shown that by sampling the slow fMRI signal at a very high speed (TR, 125 ms) and applying multivoxel pattern analysis, we were able to decode visual stimulus information represented at a time faster than the peak of the fMRI signal magnitude. This approach achieved the information readout from the fMRI signals by at least 2 s. We further demonstrated the importance of multivoxel pattern information for this result. The information in the pattern was represented at a nearly constant delay relative to the onset of neuronal activity, independent of the voxel-wise spread in fMRI signal delay and was able to be extracted from the voxels without large veins. These results indicate that our method is capable of extracting neural information representations from fMRI signals at high speed and independent of vascular dynamics. This approach can potentially be used for high temporal resolution decoding in other brain regions.

Although the primary purpose of this study was to acquire fMRI signals at high speed to evaluate how fast neural information could be read out using the slow event-related experimental design (Figure 1), fast signal acquisition was found to be useful in the functional mapping of brain activity based on a block-design experimental design as well. Even with only one run of data lasting only about 6 min, we were able to obtain a clear activity map (Figure 2). It is not obvious whether the increase in the number of data samples has a significant effect on the improvement of the statistical power because the fMRI signal has a very strong temporal autocorrelation and the fast sampling of the slow fMRI signals could merely increase the correlated or similar signals, not providing the increase of independent information. However, the previous study (11) showed that a shorter TR yields a higher statistical power than a longer TR given a fixed overall imaging time, suggesting that increasing the number of data samples would be advantageous in the statistical test even if the signals are correlated. This evidence implies that our ultra-fast sampling of the fMRI signal would also be effective for even the functional mapping of brain activity in a block-design type experiment such as used in the localizer session in this study.

In particular, the advantage of the fast sampling becomes more evident when comparing the rising phase of the magnitude of fMRI signals across voxels and when looking at the decoding performance. The fMRI signal can be modeled as the convolution of neural activity with the hemodynamic response, whose time constant is very large. Thus, a TR of 2 s would suffice to recover the time course of any fMRI signals according to the sampling theorem. However, since our purpose was not the reconstruction of the time course of the original brain activity but the identification of the latency to extract represented information, such fast sampling was useful for comparing fine differences in the time course of the magnitude of fMRI signals and decoding performance.

Although we used a short TR of 125 ms, it might be interesting to examine how much we could increase TR to see similar results. Since we applied temporal smoothing before decoding, the signal acquisition could be much slower. If TR could be increased, we might be able to scan more slices to cover a larger brain area. Such a setting would be more practical for various studies that need to analyze temporal dynamics of the fMRI signals in multiple brain areas such as inter-areal functional connectivity analyses.

The information of object category images is known to be encoded in the multivoxel pattern of fMRI signals in the ventral high-order visual cortex (12, 13), and the present study obtained the same results (Figures 2 and 3), showing the validity of our methods to capture the target signal patterns. fMRI signals for the “animal” and “vehicle” categories differed in their mean amplitude averaged in the ROI (Figure 3), but the use of such univariate information (mean amplitude only) achieved lower accuracy of the prediction, and its peak timing was later than when using the multivoxel patterns (Figure 4). These results suggest that the information we read out was indeed derived from object categories and was represented in the very early rising phase of the fMRI signal after the stimulus onset.

More interestingly, this rapid decoding had a consistent and short latency, independent of the variation in the latency of the fMRI signals used for the decoding (Figure 6). The multivoxel pattern analysis that we used, together with the fast sampling of fMRI signals, has a sensitivity to signal components containing information, independent of the spread in hemodynamic latency or magnitude that is associated with variable voxel-wise sampling of the vasculature.

Further analyses using the sparse model support this hypothesis. As Figure 7 shows, the venous voxel selectivity dropped at the timing when decoding performance increased. This result suggests that the non-venous voxels should have enough information to achieve accurate decoding performance in the early rising phase of the fMRI response. An ideal signal acquisition of fMRI experiments would avoid signal components stemming from large veins that have less spatial specificity and large temporal variation (2) and pick those from the capillary bed close to the neuronal tissues, but these components are usually mixed together. However, it is highly likely that the combination of the fast sampling and multivoxel analysis of fMRI signals provides a natural filter that eliminates large vessel signal components, using only the capillary-related components in the early rising phase that strongly reflect neuronal activity.

One might think that this argument about information retrieval from non-venous voxels counters previous studies claiming that the information source of visual decoding from fMRI signals is large veins, not stemming from functional cortical microstructure that has feature representation (14, 15). However, the previous studies used a block design with a conventional TR, and thus, their results would be influenced by the signal components with a large amplitude, which more likely stem from the large veins and evolve over time. In contrast, our method allows us to extract the early and relatively small signal components, which more likely reflect neural activity represented in non-venous regions during the short latency period. Hence, the difference in these studies might come from the difference in the time at which fMRI signals are analyzed.

In this study, we assessed decoding performance using event-related fMRI. The rapid decoding may filter out large-vessel contributions yet is still reliant on a hemodynamic effect – presumably in capillaries. The rapidity of this effect and the remarkable temporal consistency provide promise for further elucidating relative onsets of different brain regions and perhaps the cascade of sequential neuronal activity that accompanies different behaviors and cognitive processes.

However, each brain area may have subtle but systematic differences in predominant vascular architecture, yielding non-neuronal variations in decoding delay. Such intrinsic differences in local hemodynamics can be visualized by breath-holding experiments (16), which are thought to be independent of the visual cortical hierarchy and its functional connectivity.

We found that visual stimulus information can be extracted from fMRI signals within about 2-3 s after the stimulus onset; however, we also observed subtle effects as early as 1 s after the stimulus onset. While we cannot identify the fMRI signal component that is the source of this early increase in decoding accuracy, one hypothesis is that multivoxel decoding is detecting information in the elusive “initial dip” buried in the fMRI signal rising phase for some participants. Because multivoxel pattern analysis bootstraps fMRI sensitivity based on information content, it might be detecting this putative initial dip as an information source to achieve statistically significant decoding performance. Further refinement of this novel approach to fMRI data paradigm design and analysis may open up new avenues by which more rapid and subtle fMRI dynamics may be captured.

## Materials and Methods

### Participants

Fourteen healthy volunteers participated in functional and anatomical imaging sessions, and eight out of them further participated in a vein-mapping session after granting informed consent under an NIH Combined Institutional Review Board-approved protocol (NCT00001360). Data from one female subject was excluded because of reported peripheral nerve stimulation during the functional imaging session.

### Stimulus and experimental design

We had two types of functional imaging sessions to acquire fMRI signals mainly from the visual cortex: 1) decoding session, and 2) functional localizer session.

The decoding session had 8-10 runs depending on participants. Each run consisted of 32 stimulus image presentations, each of which had a duration of 500 ms followed by an inter-stimulus interval (ISI) presenting a bulls-eye fixation spot on a gray background, and extra rest periods were added at the beginning (12 s) of each run. To avoid participants predicting the onset of stimulus presentation, the duration of each ISI was randomly varied between 8.5 and 13.5 by a multiple of the duration of a single fMRI volume acquisition. The total duration of each run was always 380 s.

Stimulus images were either of eight naturalistic object categories (“male”, “female”, “Siberian husky”, “toy poodle”, “airliner”, “fighter”, “jeep”, “sports car”) sampled from ImageNet database (17). The background of the original stimulus images was removed and replaced with homogenous gray so that only the target object was presented to participants. Each stimulus image was presented at the center of the screen, and the size was adjusted to subtend 12 x 12 deg (horizontal x vertical). The same fixation spot as in the ISI period was present on the top of each stimulus image. Because all stimulus images were different exemplars, four exemplars for each category were presented for each run and thus 32-40 exemplars were presented for the whole experiment.

The functional localizer session consisted of one run with eight stimulus blocks for each participant. Each stimulus block was 20-s long presenting intact naturalistic images, followed by a 20-s period presenting scrambled naturalistic images except that the duration of the last scrambled block was 10 s. An extra scrambled stimulus block was added at the beginning for 10 s, and then the total duration of the functional localizer run was 320 s. Within each intact stimulus block, a stimulus image was flashed at a cycle of 500 ms on and 500 ms off (homogeneous gray) and another intact image was chosen for the next flash. Similarly, scrambled images were flashed with an intervening homogeneous gray image in the scrambled stimulus block. The intact stimulus images were sampled from the same categories of ImageNet as used in the decoding session and were also preprocessed similarly. The same image exemplars were not used between the decoding presentation session and the functional localizer session. The scrambled images were generated by scrambling the intact images selected for this session. The size of the presented images was also similarly scaled as in the decoding session.

In these sessions, participants viewed the stimulus sequence while maintaining fixation. To ensure alertness, they were instructed to press a button in the scanner when detecting a change in the fixation color from green to red for 250 ms at a random interval between 1 and 9 s.

### MRI acquisition

MRI data were obtained using a Siemens MAGNETOM 7T scanner located in NIH/NIMH with a 32-channel head coil. A T2*-weighted multiband (multiband factor of 3) echo-planar imaging (EPI) sequence (18) was used to acquire nine slices of functional images to cover occipital and occipitotemporal areas along the ventral pathway (TR, 125 ms; TE, 20.8 ms; flip angle, 15 deg; field of view, 192 x 192 mm; in-plane voxel size, 3 x 3 mm; slice thickness 5 mm; slice gap, 0 mm; in-plane acceleration (iPAT) factor, 2) for both decoding and functional localizer sessions.

T1-weighted magnetization-prepared 2 rapid acquisition gradient echo (MP2RAGE) (19) structural images of the whole-head were also acquired (TR, 6,000 ms; TE, 3.02 ms; TI1/TI2, 800/2700 ms; flip angle 1/2, 4/5 deg; field of view, 224 x 224 mm; in-plane voxel, 0.7 x 0.7 mm; slice number, 224, slice thickness, 0.7 mm).

A T2*-weighted gradient-echo 3D fast low angle shoot (3D-FLASH) sequence was also used to acquire a susceptibility-weighted image (SWI) to depict vein locations in an area that matched the functional images (TR, 23 ms; TE, 15 ms; flip angle, 20 deg, field of view, 165 x 220 mm; in-plane voxel size, 0.491 x 0.491 mm; slice number, 128; slice thickness, 0.5 mm).

Physiological signals (pulsation and breathing) monitoring was also performed with the MRI scanning experiment. Pulsation was monitored using an infrared sensor attached to the tip of the index finger of the hand that was not used to perform the task. Breathing was monitored by a stretch sensor rolled around the participant’s abdomen. Both signals were measured by BIOPAC (BIOPAC System, Inc.) at a sampling rate of 1 kHz.

### MRI data preprocessing

The first 8-s scans of each functional run of the decoding session and the functional localizer session were discarded to avoid contamination of instability of the MRI scanner. After removal of spike noise from fMRI signals, retrospective image correction was performed to remove physiological motion artifacts by using cardiac and respiration signals (20). The fMRI data of both sessions then underwent within-session motion correction. The whole-head high-resolution anatomical image was coregistered to an fMRI volume with the least motion in the decoding session. The coregistered anatomical image was then used as the target image to which the motion-corrected fMRI data of the functional localizer session were coregistered using an fMRI volume with the least motion in the functional localizer session as a reference image. The fMRI data of both sessions were then reinterpolated by the voxel coordinate defined by an fMRI volume with the least motion in the decoding session. All these preprocessing steps were performed in AFNI (21) using the standard “super-script” afni_proc.py (https://afni.nimh.nih.gov/) (22).

### Regions of interest

The fMRI data of the functional localizer session were used to define higher visual cortical (HVC) areas that were responsive to object images. These regions were served as the region of interest (ROI). Linear regressors were used to define the time during which intact object images and scrambled object images were presented, and their coefficients were computed by a generalized linear model with a restricted maximum likelihood method. Statistical contrast of the coefficients between intact and scrambled object-images regressors was then computed by t-test for each voxel. fMRI volumes were not taken into account for these computations if the Euclidean norm of the motion alignment parameter relative to the previous volume was larger than 0.3 or if a large fraction (10%) of voxels were classified as outliers that showed excessive deviation from the time series of the voxel. The HVC areas were then defined by voxels showing uncorrected p-values larger than 0.001 with a cluster size larger than 40 voxels. Voxels outside the individual gray matter were excluded. These procedures were done for each individual participant separately.

### Time-resolved decoding

The fMRI data underwent signal preprocessing in the following order. First, the voxels within the ROI were selected according to their statistical significance for the functional localizer session. The number of the selected voxels was 400 in the descending order of the t-value unless otherwise stated. For each run, outlier voxels were identified as those with amplitude exceeding +/-100% of signal change relative to the mean amplitude of the run. Such voxels were removed from further analyses. The time course of the fMRI signal of each voxel underwent linear trend removal within each run, and its amplitude was normalized relative to the mean amplitude of the first 4-s rest period in each run to minimize the baseline difference across runs.

A linear support vector machine (SVM) (23), or decoder, was then trained with the preprocessed fMRI data. We evaluated the decoding performance of the presented stimulus categories (only binary classification of “animal” (consisting of “male”, “female”, “Siberian husky”, and “toy poodle”) vs. “vehicle” (consisting of “airliner”, “fighter”, “jeep”, and “sports car”) was performed) by using a leave-one-run-out cross-validation procedure. The training and testing of the decoder were performed at each volume (i.e., each 125 ms) after the stimulus presentation and the timing was shifted to evaluate all time points in each trial, providing the time course of decoding performance in the trial. To suppress fluctuations from high-frequency noisy components, the preprocessed fMRI data were temporally averaged within ±3 volumes around the specified target volume for the decoding (i.e., fMRI signals from 7 volumes are temporally averaged and used to define a new smoothed data point at the target volume) before training and testing the decoder.

### Time-resolved sparse decoding

To identify informative voxels to extract presented stimulus information, we further applied a sparse decoding technique to the fMRI data in the same way. Sparse logistic regression (SLR) decoder (7) was used as a sparse decoder for this analysis. SLR is based on a hierarchical Bayesian model using automatic relevance determination (ARD) prior, which promotes sparse selection of voxels and assigns non-zero weights for only a small number of voxels. The remaining voxels have zero weights and thereby do not contribute to the decoding. The selected voxels can be considered as “informative” voxels to extract the presented stimulus information. We run this decoding in a time-resolved manner as explained in the previous methods using SVM and identified the temporal changes of the informative voxel sets.

### Analysis of venous voxel locations

To examine whether the informative voxels identified by the sparse decoding analysis include large vasculature, we identified venous voxels using T2-contrast-enhanced susceptibility-weighted image (SWI). The SWI was acquired at much higher spatial resolution (0.491 x 0.491 x 0.5 mm) to identify venous voxels as precisely as possible. Since the vein locations can be illustrated as a dark region, the venous voxels were defined as those with a lower intensity than a threshold value. To define the venous voxel map as fine as we can, three steps were applied as Geißler et al. (2013)(24): 1) make an initial binary map by defining the first threshold values, 2) smooth the binary map, and 3) make the fine venous binary map by defining the second threshold values. To define the first threshold value, we calculated a histogram of voxel intensity values for each individual SWI. We found that the histogram typically showed two distinct peaks, with the distribution splitting around pixel values between 100 and 200. We then adopted 150 as the first threshold. Then the map was smoothed using the “smoothn” MATLAB function with the smoothing parameter S of 1. Lastly, the second threshold was defined by visual inspection, resulting in varying 0.05-0.15 across participants. Each SWI voxel was labeled as either venous or non-venous according to these procedures. Because a functional image voxel was much larger than an SWI voxel, we classified each functional image voxel as venous or non-venous by calculating the proportion of venous SWI voxels it contained. In this study, we identified a functional image voxel as a venous voxel if this proportion exceeded 50%. Then we calculated how many voxels were venous out of the sparse-decoder-selected voxels (“venous voxel selection rate”) for each volume, which generates a time course of relative contribution of venous/non-venous voxels for the prediction of the presented stimulus information. Because the absolute values of the venous voxel selection rate varied widely across participants, we normalized each participant’s time course by converting it to z-scores as

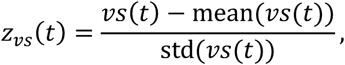

where *Z*_*vs*_(*t*) is the z-score of the venous voxel selection rate at time *t*, *vs*(*t*) is the original venous selection rate at time *t*, mean(*vs*(*t*)) and std(*vs*(*t*)) are the mean and the standard deviation of the venous selection rate over the single-trial time course, respectively. The resulting time courses of the z-scores were then averaged across participants.

## Acknowledgments

We thank Martin Hebart for discussions on experimental design and results, Chung Kan for technical assistance with 7T MRI scanning, and Paul Taylor and Rick Reynolds for technical assistance with fMRI signal preprocessing using AFNI. This work is partially supported by JSPS KAKENHI (JP20H00600, JP18KK0311, JP17H01755, JP25H01138), JST PRESTO (JPMJPR1778), and Yamada Science Foundation. Data used in the preparation of this work were obtained from the NIMH’s Section on Functional Imaging Methods (SFIM; Principal Investigator: Peter A Bandettini). This research was supported in part by the Intramural Research Program (ZIAMH002783, ZICMH002888) of the National Institutes of Health (NIH). Clinical Protocol: NCT00001360. The contributions of the NIH authors were made as part of their official duties as NIH federal employees, are in compliance with agency policy requirements, and are considered Works of the United States Government. However, the findings and conclusions presented in this paper are those of the author(s) and do not necessarily reflect the views of the NIH or the U.S. Department of Health and Human Services.

