## Supplementary Figure 1 for "Rapid decoding of neural information representation from ultra-fast functional magnetic resonance imaging signals"

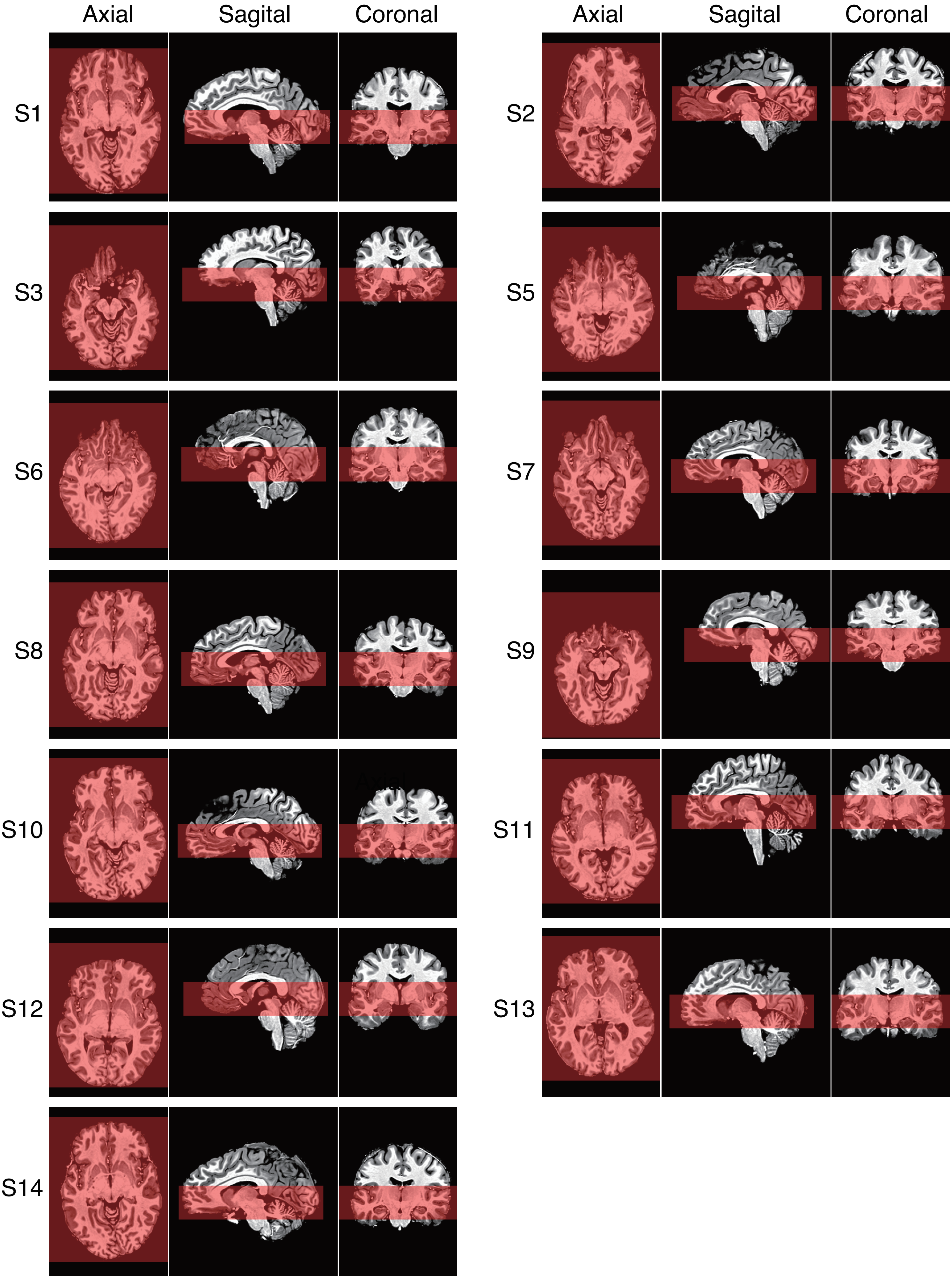


**Supplementary Figure 1: The field of view of functional image scans.**
Red transparent regions show the spatial coverage (field of view) of functional image scans overlaid on the anatomical image for each participant. The displayed sections for each axis are aligned to the center of the scanned volume.
