## Supplementary Figure 2 for "Rapid decoding of neural information representation from ultra-fast functional magnetic resonance imaging signals"

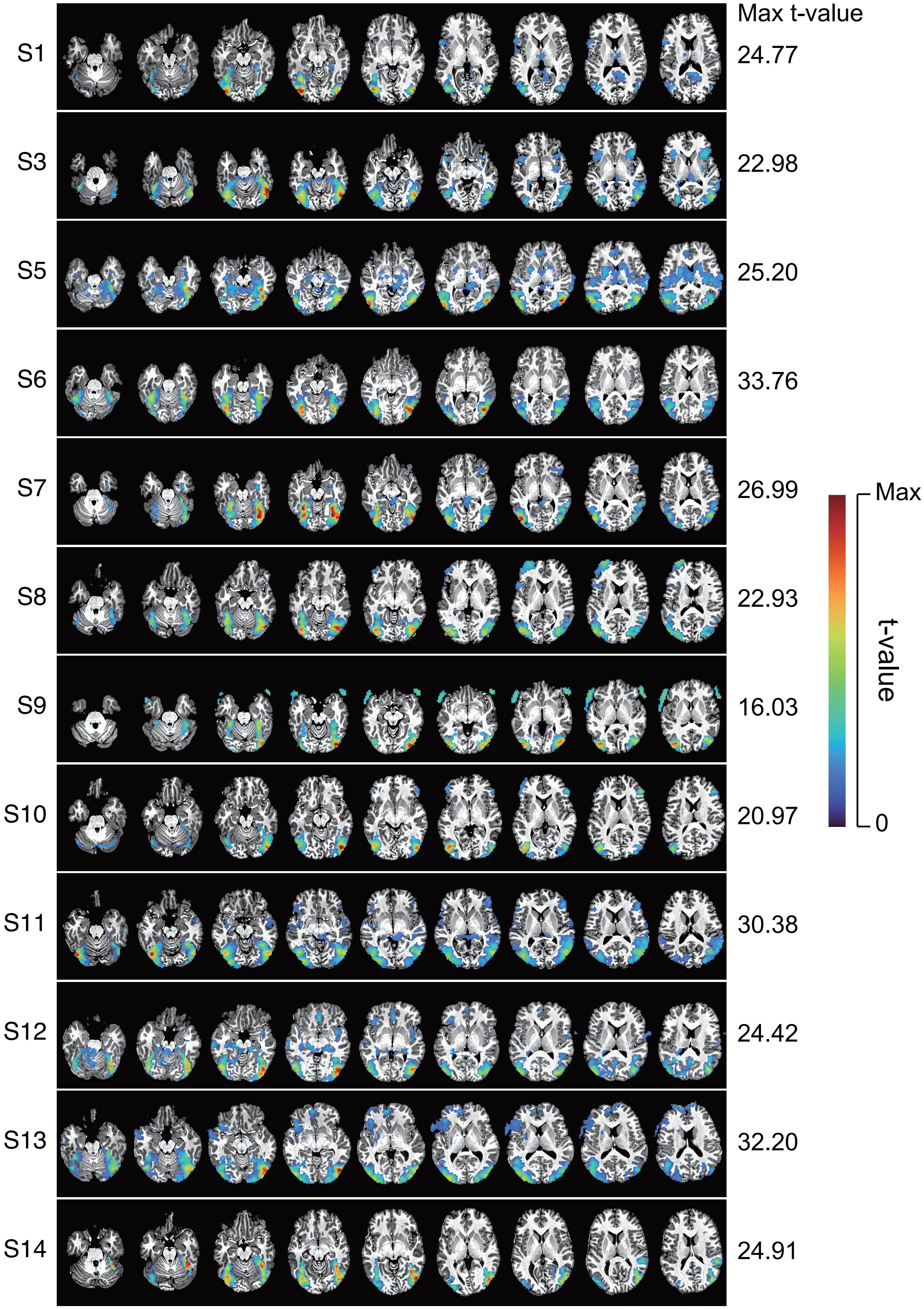


**Supplementary Figure 2: Functional localization with short TR signal acquisition for all participants (except S2).** Statistical t-value maps of fMRI signal contrast between intact – scramble object images for each functional slice are shown on the anatomical image. The in-plane voxel size is 3 x 3 mm and the slice thickness is 5 mm (see also Methods; each image is 5-mm apart in the inferior-superior direction). Voxels are colored according to the same criteria as Figure 2A for each participant (the maximum t-values are displayed to the right of the each participant result).
