## Supplementary Figure 3 for "Rapid decoding of neural information representation from ultra-fast functional magnetic resonance imaging signals"

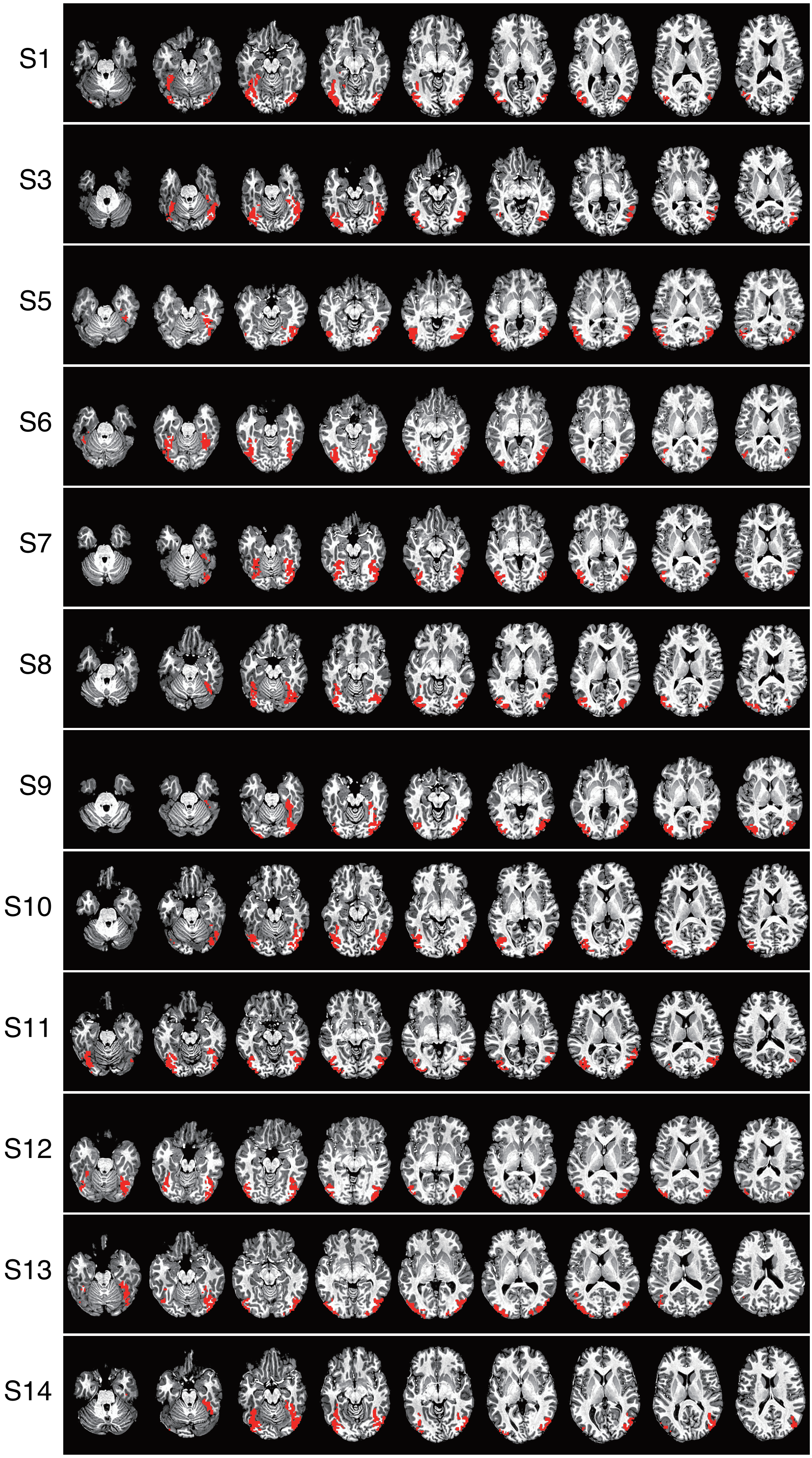


**Supplementary Figure 3: Voxel masks for input to the decoding analysis (except S2).**
The location of the top 400 t-value voxels from the individual gray matter was displayed in the same way as Figure 2B.
