## Supplementary Figure 4 for "Rapid decoding of neural information representation from ultra-fast functional magnetic resonance imaging signals"

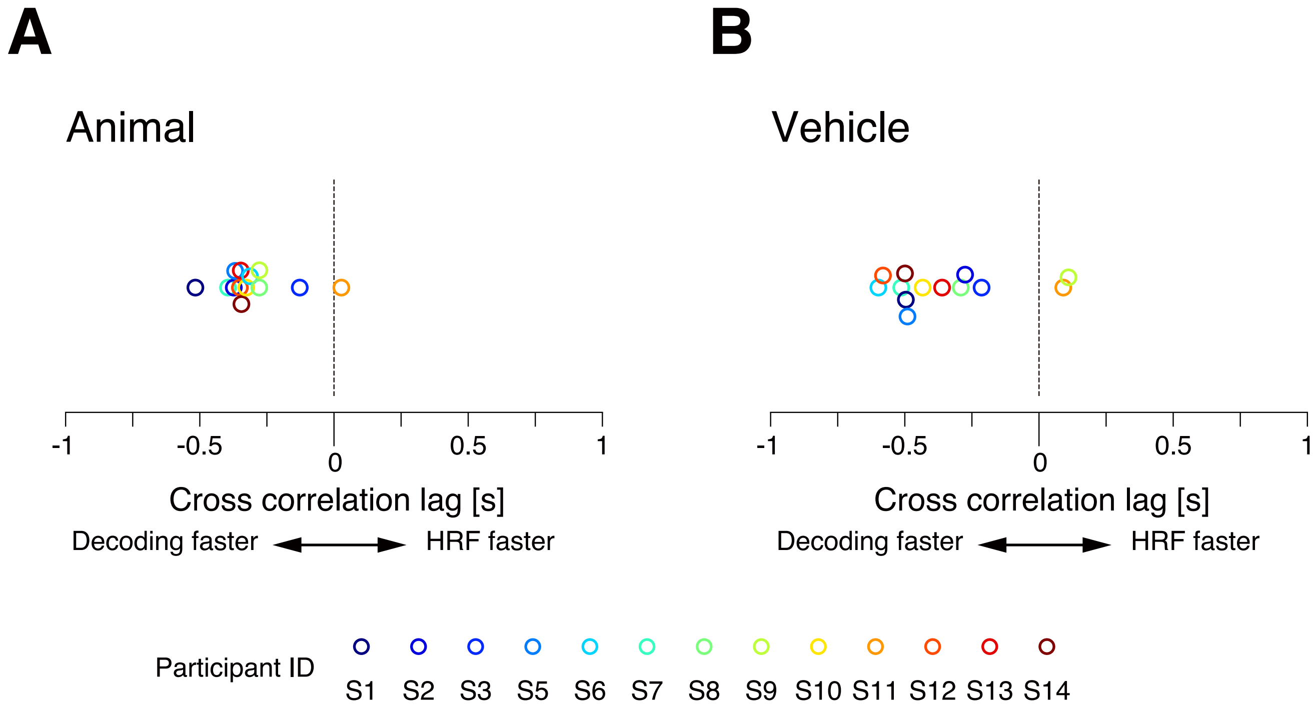


**Supplementary Figure 4: Individual variations in the lag between the time course of decoding performance and HRFs.**

The mean cross-correlation lag between the time course of decoding performance and single voxel HRFs for all participants for “animal” (A) and “vehicle” (B) categories. Unique colors are assigned to each participant for both panels. Dashed lines indicate the zero lag. Markers are jittered vertically to avoid overlap for visualization purposes.
